# Region-specific auditory and visual plasticity in mouse visual cortex after early visual deprivation

**DOI:** 10.1101/2024.08.31.610645

**Authors:** Huub Terra, Matúš Halák, Leander de Kraker, Christiaan N. Levelt

## Abstract

A lack of early visual experience causes pronounced lasting auditory and visual cross-modal changes. However, the region-specific cross-modal organization of the visual cortex at the single-neuron level remains unknown. Here, we used two-photon calcium imaging in awake mice that were reared in darkness from birth and were consequently exposed to light in adulthood to map auditory and visual responsiveness of single neurons. We targeted neurons in the primary visual cortex (V1) and higher visual cortical areas (HVAs) that resemble the ventral and dorsal stream regions. We found that lateral dorsal stream areas showed a pronounced increase in auditory response strength, even after accounting for tone-induced whisker movement. Strikingly, this was accompanied by a decreased visual drive, measured by the number of recruited neurons, response strength, population orientation decoding accuracy and neural manifold size, although these visual effects were more widespread across cortical regions. Together, these results provide a comprehensive functional map of auditory and visual cross-modal changes after a lack of early visual experience across the mouse visual cortex. Moreover, our results suggest that a lack of visual drive of dorsal stream regions might provide an opportunity for the remaining senses to take over.

## INTRODUCTION

Our visual cortex is wired to predominantly process visual input from our eyes. However, in congenitally blind individuals, the deprived visual cortex can shift its processing to one of the remaining sensory modalities, such as hearing, through a process called cross-modal plasticity^1–4^. In humans, this audio-visual cross-modal plasticity is accompanied by an increased ability to process sounds, for example improved pitch discrimination^5^ or auditory localization^3^, possibly due to enhanced processing of auditory information by the visual cortex. At the same time, visual processing is impaired but still possible, as seen after restoration of sight in adulthood^6–8^. After adult reversal of visual impairments, sound-evoked activity in the visual cortex can remain elevated^6,7,9^. This suggests that the cross-modal rewiring after early visual deprivation might remain fixed into adulthood and thereby prevent full recovery of vision. How this is organized across the visual cortex at the single-neuron level remains unknown.

The visual cortex contains specialized regions that process different types of sensory information: the dorsal stream regions process movement-related information, the ventral stream regions object-like information, and the primary visual cortex (V1) processes simpler features. In humans, these regions have been suggested to be differentially sensitive to early visual experience, with the dorsal stream regions in particular appearing underdeveloped after loss of early visual experience^10^. In rodents, effects of early visual deprivation on visual processing have also been reported primarily in V1^11,13,15,20^. Dark rearing mice from an early age resulted in lower responsiveness of populations of neurons in dorsal stream-like areas to visual stimuli^12^. This shows that dorsal stream regions are more vulnerable to a lack of visual experience than V1 or ventral stream regions. However, whether this also means that single neurons of the dorsal stream regions become more responsive to sounds remains unknown.

Animal models, and especially mouse models with their abundant options for cell targeting and manipulation, are essential to understand the cell-type-specific changes underlying cross-modal changes after loss of early vision. Cellular changes ranging from genetic to functional have been reported in the visual cortex after loss of early vision^14,16,17^. Layer 2/3 and layer 4 cells in V1 change their genetic identity after dark rearing from an early age^16^. Complete removal of retinal input to V1 by binocular enucleation even shifts the identity of V1 cell types towards those of higher visual cortical regions. Together, this suggests that visual experience is crucial for the formation of the sensory identity of visual cortical regions^17^. In terms of visual processing, early visual experience is essential for the sharpening of basic properties such as orientation and direction tuning^18–20^, although direction tuning in the primary visual cortex of mice has been suggested to be independent of early visual experience^19^. Overall, the neural population is organized is such a way that it can optimally detect stimuli, such that each stimulus (like grating or direction or orientation) is well separated in a so called sensory manifold^21,22,24^. In terms of auditory processing, single neurons of the visual cortex respond sparsely to sound, being either activated or suppressed by tones^23^, and contribute to audio-visual multisensory integration^25–30^. In animal models, loss of early visual experience can cause visual cortical neurons to become more responsive to sounds^31–33^. While most of these changes in responsiveness have been reported in V1, increased auditory activation of lateral extrastriate area V2 has also been reported^32^. However, whether visual experience-dependent changes in auditory responses differ between visual cortical subregions remains unknown. A complicating factor is that recent studies have shown that sound-induced neural activity is often caused indirectly by sound-induced facial movement^34,35^. Thus, a comprehensive study of movement-controlled auditory and visual responses of region-specific visual cortical single neurons and populations neurons after loss of early visual experience is missing. Additionally, whether these responses are reversible in adulthood by introduction of vision remains unknown.

We therefore set out to explore the effect of loss of early visual experience on auditory and visual processing throughout V1 and the higher visual cortical areas (HVAs) anteromedial (AM), posteromedial (PM), anterior (A), rostrolateral (RL), anterolateral (AL) and lateromedial (LM). We hypothesized that dorsal stream areas^36–38^ AM, PM, A, RL and AL would be particularly sensitive to a lack of visual experience, showing increased auditory and decreased visual processing. Visual experience was manipulated by dark rearing mice from birth. In awake, head-fixed mice that were free to run, functional auditory and visual changes at the cellular level were recorded using calcium imaging under a two-photon (2P) microscope in response to pure-tone and moving-grating stimuli. Recordings were targeted to V1 and the combined HVAs: AM/PM (medial dorsal stream), A/RL/AL (lateral dorsal stream) and LM (ventral stream). For auditory stimuli, we corrected for the sound-induced arousal effect using video monitoring and face movement analysis. We found that a loss of early visual experience has a pronounced effect on single neurons in HVAs, especially the dorsal stream regions, with an increase in auditory processing and a decrease in visual processing capacity on both a single neuron as population level. This study provides the first comprehensive mapping of functional auditory and visual cross-modal changes after loss of early visual experience and can contribute to a cellular-level understanding of the cross-modal changes after early loss of vision.

## RESULTS

### Dark rearing increases responsiveness of neurons in anterior higher visual cortical areas to pure tones

To test the effect of loss of early visual experience on auditory processing of single neurons across visual cortical regions, we presented pure-tone stimuli to naive mice. One group of mice was reared in absolute darkness from birth and therefore lacked any visual experience, and a second group was reared in normal light/dark conditions (**Fig. 1A**). Adult mice were first placed under a wide-field macroscope to determine the location of visual cortical areas using population receptive field mapping and the Allen Brain reference atlas (**Fig. 1B**)^37,39,40^. Activity of single layer 2/3 neurons was recorded using 2P calcium imaging in awake, head-fixed mice that were free to run while pure tones were presented from a single speaker in front of the mouse (**Fig. 1C**). Stimuli were presented in darkness while facial movement and run speed were recorded under infrared light.

**Figure 1.**
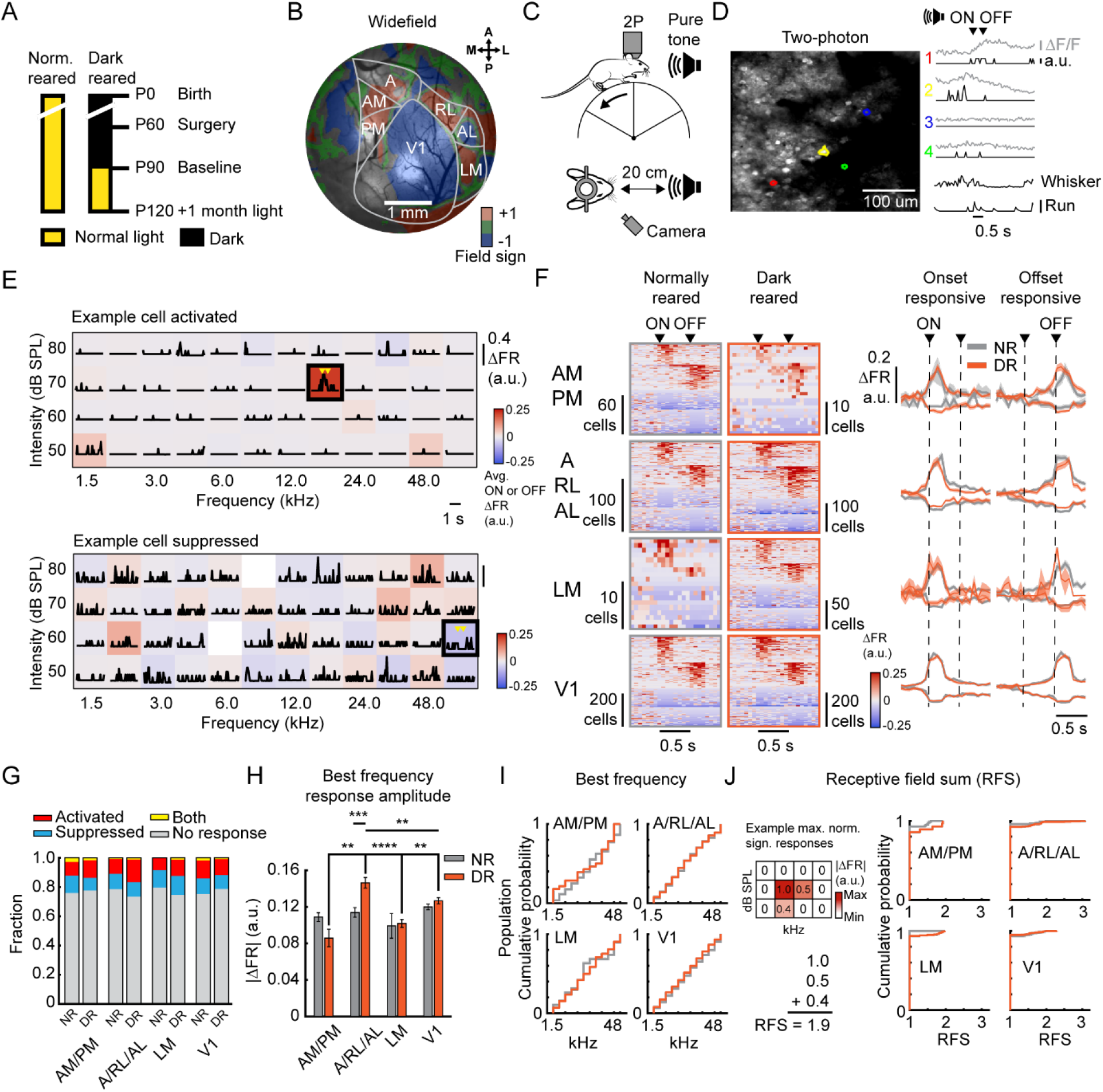
Dark rearing increases responsiveness of neurons in anterior higher visual cortical areas to pure tones. (A) Experimental groups and timeline. (B) Example widefield epifluorescence microscopy field of view overlayed with a field sign map and the Allen Brain institute reference atlas. (C) Schematic of the 2P microscope experimental setup for presentation of pure tone stimuli. Tones are presented for 0.5 s at 12 frequencies ranging from 1.5 kHz to 48 kHz and half-an-octave step with half-an-octave step in between and at 4 intensity levels between 50 and 80 dB sound pressure level (SPL) with 10 dB SPL steps with 2-4 s in between stimuli. Facial movement is recorded using an infrared camera under infrared background light. (D) Left, example 2P microscopy field of view of the visual cortex with GCaMP8m-expressing cells. Right, Example responses of four cells to a pure tone sound (12 kHz and half-an-octave step at 70 dB SPL). Grey traces, calcium fluorescence (ΔF/F), black traces below grey traces, deconvolved spikes (a.u., bar = 1 a.u.), black traces, whisker energy and run speed (bar = 0.1 cm/s), arrows indicate tone onset and offset. (E) Example frequency response areas of a cell excited (top) and inhibited (bottom) by pure tones. Traces indicated baseline-corrected mean response over trials in a.u (ΔFR). Raster plot, trial-averaged ΔFR during 0-0.2s after tone onset or offset. Empty slots indicate removed tones due to having less than 8 trials left over after removal of whisker-movement trials. Box indicates strongest significant response to a tone, i.e., best frequency. (F) Significant responses to pure tones. Per cell the best frequency response is displayed. Left, Plot sorted on response amplitude and response category (tone onset or offset). Right, mean responses over cells that respond strongest with excitation (top), inhibition (bottom), to tone onset (left) and offset (right), (G) Fraction of response types. Fisher’s exact test, p > 0.05. (H) Response amplitude (absolute ΔFR) at the best frequency for tone-responsive neurons. Two-way repeated measures ANOVA with Tukey’s multiple comparison’s test between groups and within groups, Rearing, F(1, 1912) = 0.05311, p = 0.04662, Cortical region, F(3, 1912) = 6.582, p < 0.0002, Interaction, F(3, 1912) = 4.702, p < 0.0028, n = 19 – 550 cells per group from n = 3-6 mice per group. (I) Cumulative probability function of best frequencies over all tone-responsive neurons. Mann-Whitney U test, p > 0.05. (J) Receptive field sum (RFS). Left, illustrative schematic of RFS calculation. Right, Cumulative probability function of the RFS of all tone-responsive neurons. Mann-Whitney U test, p > 0.05.

Recent work showed that sound-induced activity of V1 neurons is predominantly correlated with facial movement rather than with sound itself^34,35,41^. Movement of the whisker pad in particular was highly predictive of neuronal activity^35^. Temporally, the motor-related component of neural activity was strongest after about 200 to 300 ms, whereas the auditory component was strongest before that time^34^. As expected, we found that pure tones elicited whisker movement (**Fig. S1A and B**). Both tone onset and offset evoked whisker movement (**Fig. S1B and D**). Normally reared and dark reared animals showed frequency- and intensity-dependent whisker movement (**Fig. S1C**). Tones of higher intensity (**Fig. S1D**) and near the peak sensitivity of mouse hearing, around 6 kHz^42^ (**Fig. S1E**) induced the strongest whisker movement. No effect of rearing condition was observed. To minimize arousal-related components in tone-induced neuronal activity, we 1) removed all trials with Z-scored whisker movement energy above two, 2) improved the temporal resolution of our fluorescence signal by estimating spike times with a spike deconvolution algorithm^43^ and 3) restricted our analysis to approximately the first 250 ms after tone onset or offset.

We recorded the frequency response areas (FRAs) of neurons and observed that pure tones can activate and inhibit neurons in the visual cortex (**Fig. 1E**). Both tone onset and offset drove changes in neuronal activity in normally reared and dark reared animals throughout the visual cortical regions AM/PM, A/RL/AL, LM and V1 (**Fig. 1F**). Neurons in all cortical regions and in both rearing conditions showed similar fractions of tone-induced response types (activation, inhibition, both activation and inhibition or no response) in their FRA (**Fig. 1G**). However, dark rearing resulted in a higher tone-induced response amplitude of neurons in the A/RL/AL region at their best frequency (BF, tone with strongest response), but not in other regions (**Fig. 1H**). Further analysis showed no effect of rearing or cortical region on the distribution of best frequencies (**Fig. 1I**) or on frequency selectivity, measured as receptive field sum (RFS)^44^ (**Fig. 1J**). Together, this shows that a loss of early visual experience by dark rearing specifically increases the response strength of single neurons to pure tones in lateral dorsal stream HVAs.

### Dark rearing decreases responsiveness of neurons in anterior higher visual cortical areas and V1 to drifting gratings

The specific increase of auditory processing in A/RL/AL visual cortical regions after loss of early visual experience could suggest that this region is especially vulnerable to a lack of early visual experience. We therefore tested whether single-neuron responses to visual stimuli would correspondingly decrease after dark rearing. To this end, the same normally reared and dark reared mice were also shown full-field drifting gratings in different directions and contrasts (**Fig. 2A**) while neuronal activity was measured using 2P calcium imaging (**Fig. 2B**). As expected, we observed neurons responding to drifting gratings, often with a clear preference for a certain direction and contrast (**Fig. 2C**). Both normally reared and dark reared animals had neurons with responses to drifting gratings across the visual cortical regions (**Fig. 2D**), mainly of the activated type (**Fig. 2E**). In the A/RL/AL, LM and V1 regions, there was a decrease in the overall fraction of responsive neurons after dark rearing (**Fig. 2E**). Additionally, we observed a decrease in the response amplitude to the preferred stimulus after dark rearing in both the AM/PM and A/RL/AL regions (**Fig. 2F**). Post hoc two-way ANOVAs with Bonferroni correction targeted to A/RL/AL vs LM and A/RL/AL vs V1 showed that the decreased response amplitude was greater for A/RL/AL compared to V1 (rearing x area, F_(1, 2918)_ = 5.529, p = 0.0376) but not for A/RL/AL compared to LM (rearing x area, F_(1, 1161)_ = 2.613, p = 0.2126). Similar analyses for AM/PM vs LM (rearing x area, F_(1, 815)_ = 3.120, p = 0.1574) and AM/PM vs V1 (rearing x area, F_(1, 2572)_ = 4.456, p = 0.0698) were non-significant.

**Figure 2.**
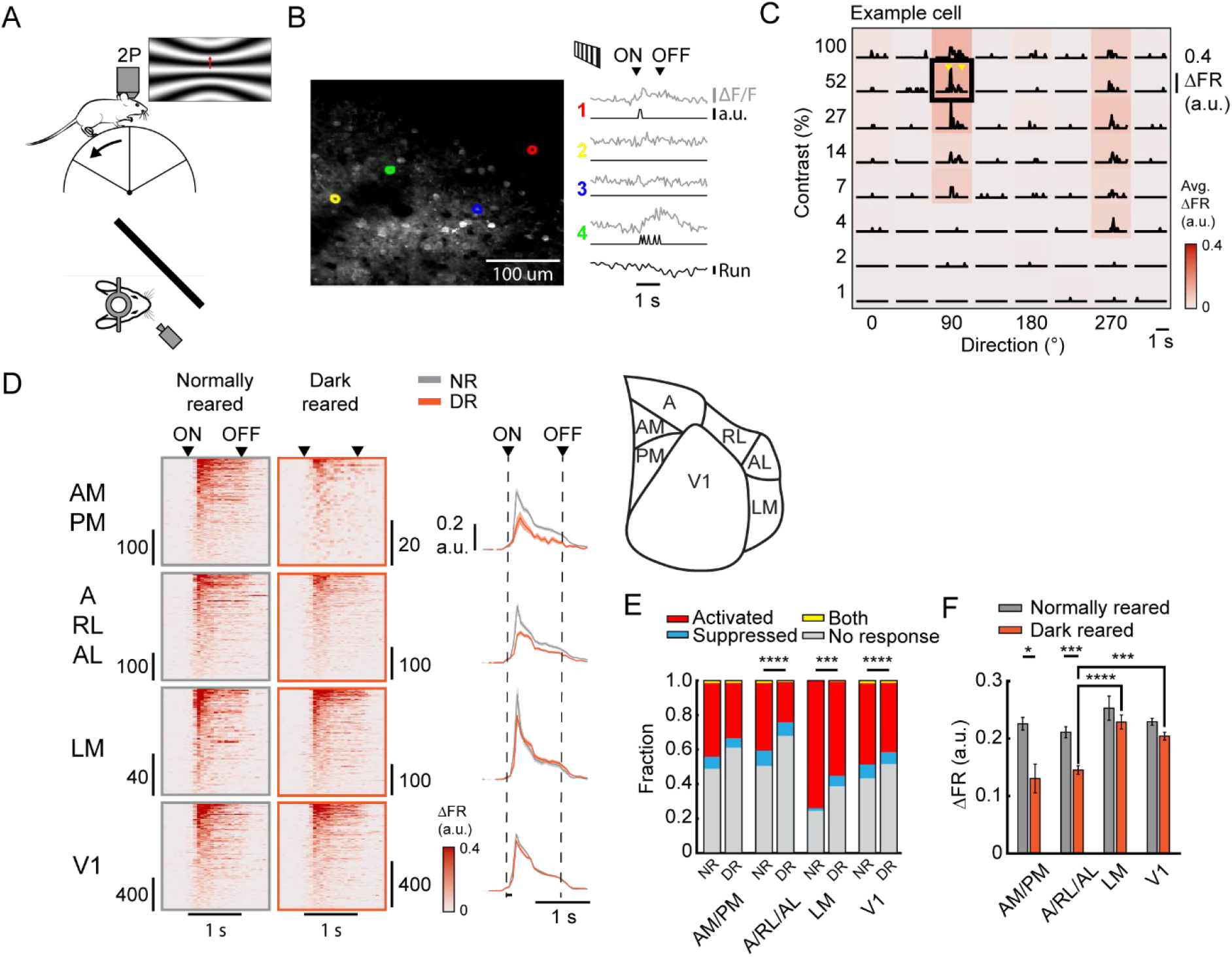
Dark rearing decreases responsiveness of anterior higher visual cortical and V1 neurons to drifting gratings. (A) Schematic of the 2P microscopy experimental setup for presentation of moving grating stimuli. Gratings are presented for 1 s at 8 directions with 45° in between and at 8 contrast levels between 1 and 100% contrast with 3 s in between stimuli and 15 repetitions per stimuli. (B) Left, example 2P microscopy field of view of the visual cortex with GCaMP8m-expressing cells. Right, Example responses of four cells to a drifting grating. Grey traces, calcium fluorescence (ΔF/F), black traces below grey traces, deconvolved spikes (a.u., bar = 1 a.u.), black traces, run speed (bar = 0.1 cm/s), arrows indicate tone onset and offset. (C) Lines, example trial-averaged ΔFR to drifting gratings. colormap, trial-averaged ΔFR during 0-0.5 s after stimulus onset. (D) Activated cells to moving gratings. Per cell the strongest response is displayed. Left, Plot sorted on response amplitude. Right, mean responses over cells, (E) Fraction of response types. Fisher’s exact test, NR x DR, AM/PM, p = 0.5490, A/RL/AL, p = 0.0001, LM, p = 0.0001, V1, p = 0.0001. (F) Response amplitude (ΔFR) at the preferred stimulus for drifting-grating responsive neurons Two-way repeated measures ANOVA with Tukey’s multiple comparison’s test between groups and within groups, Rearing, F(1, 3733) = 24.36, p = 0.0001, Cortical region, F(3, 3733) = 10.41, p < 0.0001, Interaction, F(3, 3733) = 3.085, p < 0.0262, n = 19 – 550 cells per group from n = 3-6 mice per group.

In conclusion, a lack of early visual experience reduced the responsiveness of the visual cortex to drifting gratings. While the fraction of responsive neurons decreased in V1 and HVAs, only dorsal stream regions showed reduced response amplitude.

### Dark rearing impairs orientation decoding in A/AL/RL and compresses the orientation manifold

Because dark rearing reduced the responsiveness of single neurons to drifting gratings, we next asked whether it also reduces the population decoding accuracy and collapses the neural state space, or manifold, for grating orientation and direction, and whether these changes remain plastic in adulthood.

We first decoded grating orientation and direction from population neural activity in each visual area (**Fig. 3A**). In normally reared mice, both orientation and direction could be decoded in all areas. In A/RL/AL, orientation decoding accuracy was lower in dark-reared than in normally reared mice at baseline (p = 0.0155). In contrast, orientation decoding did not differ between groups in AM/PM, LM or V1, and direction decoding did not differ in any area. After one month of light re-exposure, orientation decoding in A/RL/AL no longer differed between dark-reared and normally reared mice. Together, this shows that especially A/RL/AL neurons are dependent on visual experience for orientation, but not direction, representation.

**Figure 3.**
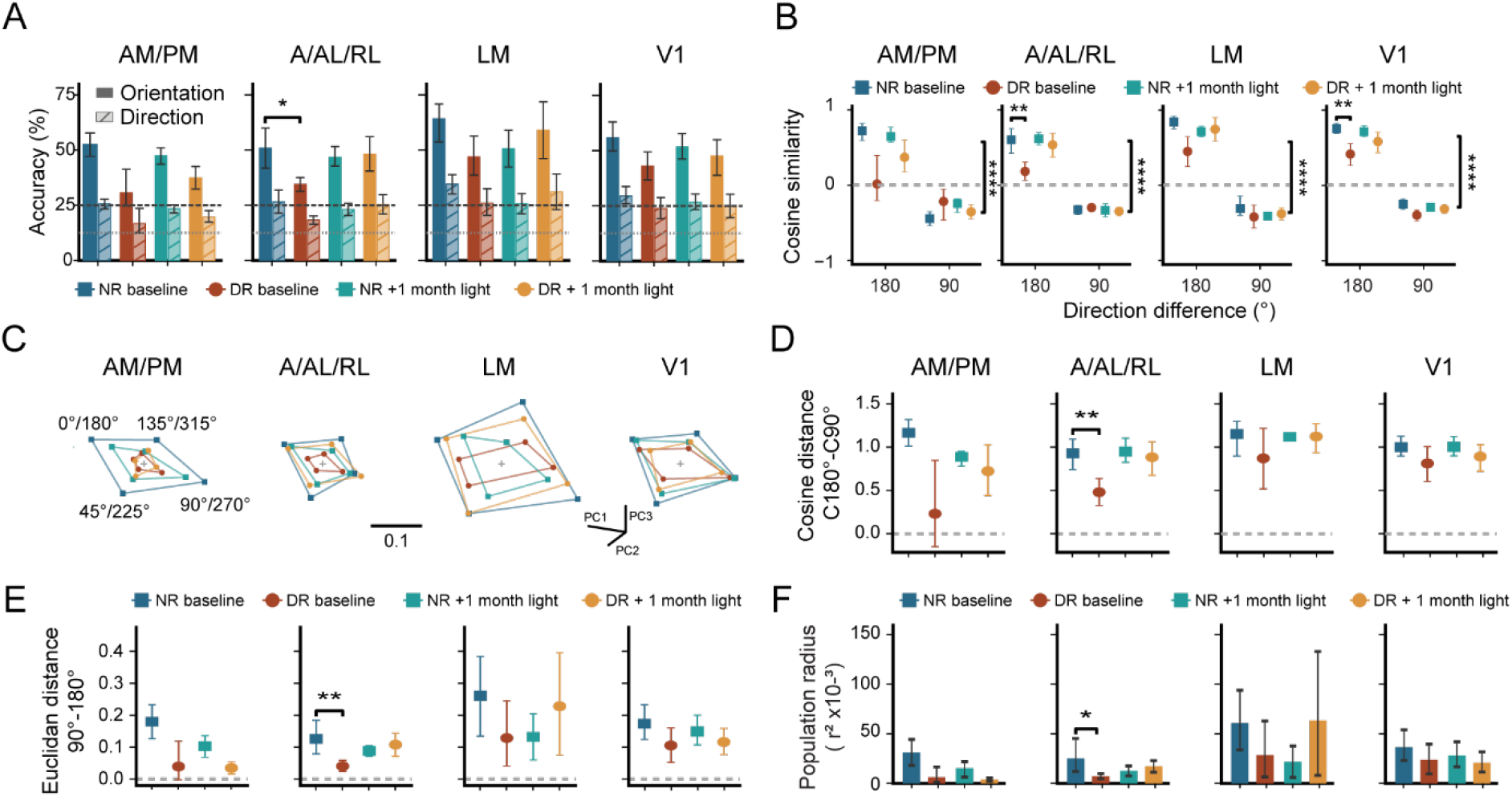
Dark rearing impairs orientation decoding in A/AL/RL and compresses the orientation manifold. (A) Cross-validated linear discriminant analysis (LDA) decoding accuracy for grating orientation (four orientations; solid bars) and direction (eight directions; hatched bars) in AM/PM, A/AL/RL, LM and V1. Dashed and dotted lines mark chance accuracy for orientation (25%) and direction (12.5%). Orientation decoding was lower in dark-reared than in normally reared mice in A/AL/RL at baseline (p = 0.0155). ANOVA-type statistics: rearing, ATS(1) = 18.59, p < 0.0001; decoder task, ATS(1) = 235.91, p < 0.0001. (B) Cosine similarity between centered population-response centroids for grating directions 180° apart (opposite directions, same orientation) and 90° apart (orthogonal orientations). Dark rearing reduced the similarity of responses to opposite directions in V1 (p = 0.00827) and A/AL/RL (p = 0.00267) at baseline. ANOVA-type statistics: rearing, ATS(1) = 19.28, p < 0.0001; direction difference, ATS(1) = 18,960.94, p < 0.0001. (C) Population responses to the four orientations, shown in a shared three-dimensional principal component (PCA) projection of the aligned population responses. Each point is the average of the two opposite-direction centroids for one orientation. Lines connect the orientations, forming one polygon per condition. Scale bar is in PCA-coordinate units. The projection is for illustration only; panels D–F give the quantitative measures. (D) Difference in cosine similarity between opposite and orthogonal directions, C(180°) − C(90°). NR vs DR, A/AL/RL baseline: p = 0.00507. (E) Difference in Euclidean distance between orthogonal and opposite directions, d(90°) − d(180°), computed in the full 10-neuron space. NR vs DR, A/AL/RL baseline: p = 0.00267. (F) Squared population radius, r²: the mean squared distance of the eight direction centroids from their center, computed in the full 10-neuron space. NR vs DR, A/AL/RL baseline: p = 0.0398. All panels use deconvolved responses averaged over 0–0.5 s after stimulus onset, pooled over the two highest contrasts (51.79% and 100%). Analyses used 1,000 random subsamples of 10 neurons per session and area. In A, B and D–F, bars and points show means in which each mouse contributes equally, after averaging across sessions within each mouse. Error bars show 95% confidence intervals from bootstrapping across mice. Colors: blue, NR baseline; teal, NR + 1 month light; red, DR baseline; orange, DR + 1 month light. Squares denote NR mice and circles DR mice. *p < 0.05, **p < 0.01, ****p < 0.0001. Mouse numbers (NR/DR, baseline; +1 month light): AM/PM: 5/4; 3/4. A/AL/RL: 8/13; 4/10. LM: 3/6; 2/4. V1: 9/10; 8/6.

To understand what changes in population coding might underlie this impaired orientation decoding accuracy, we examined the geometry of the neural state space for orientations. We first asked whether population responses are organized mainly by direction or by orientation. In every area and condition, the angle between the neural state for gratings 180° apart was more similar than the angle between the neural states to gratings 90° apart (**Fig. 3B**). This indicates that the neural population state space is more organized by grating orientation than direction.

Dark rearing reduced the similarity between responses to opposite directions in A/RL/AL (p = 0.00267) and V1 (p = 0.00827; **Fig. 3B**), suggesting a less organized neural state space. We then further tested whether dark rearing collapses the neural state space for orientations and if adult exposure to one month of light would recover this (**Fig. 3C**). We used three complementary measures: the difference in cosine similarity between opposite and orthogonal grating pairs (**Fig. 3D**); the difference in Euclidean distance between 90° and 180° gratings (**Fig. 3E**); the radius of the complete state space (**Fig. 3F**). On all three measures, the orientation state space was compressed and less organized in dark-reared compared with normally reared mice in A/RL/AL, but not in AM/PM, LM or V1. After one month of adult light exposure, the two groups no longer differed on any measure.

Together, these results show that dark rearing selectively impairs orientation coding in A/RL/AL and compresses the neural manifold for grating orientation. Both differences were no longer present after one month of light exposure in adulthood, suggesting that the adult from remains plastic.

## DISCUSSION

Here we describe cortical region-specific effects of a lack of early visual experience on auditory and visual processing at the neural population and single-neuron levels. A lack of early visual experience increased sound-evoked response strength specifically in neurons of the dorsal stream areas A/RL/AL. This dorsal stream-specific effect was accompanied by a pronounced decrease in the processing of visual stimuli by neurons in A/RL/AL as well, although the fraction of responsive neurons was lower in V1 and LM as well.

In both humans and mice, dorsal stream regions and associated functions are vulnerable during early development^10,12,45^. In humans, developmental disorders such as Williams syndrome, autism spectrum disorder and cerebral palsy are associated with impaired performance on tasks involving visual motion processing^10^. Individuals with early onset blindness also show pronounced deficits in motion processing regions such as middle temporal gyrus, middle occipital gyrus and right cuneus^3,46^. In mice, populations of neurons in dorsal stream regions, especially AM/PM and AL, develop visual tuning later after eye opening and become less responsive to visual stimuli after dark rearing compared to ventral stream regions^12^. This has led to the hypothesis that V1 input to dorsal stream regions compared to ventral stream regions is more dependent on early visual experience^12^. We expand on this work by showing that fewer neurons in dorsal stream regions AM/PM and A/RL/AL are recruited by drifting gratings and become less responsive to visual stimuli after dark rearing. Importantly, we find that these regions also become more responsive to auditory stimuli. Our results therefore provide support for the ‘dorsal vulnerability hypothesis’^12,49^ and further suggest that the reduced visual input provides an opportunity for auditory inputs to dorsal regions to strengthen.

Recently, sound-evoked activity in the visual cortex has been suggested to be caused indirectly by facial movement instead^47^. Although we indeed found a strong influence of sound on facial movement, we still observed sound-evoked activity in single neurons after removing sound-induced whisker movement trials and restricting our analysis windows to periods where purer auditory activity has been observed. Our results thus support the idea that visual cortical neurons can be activated by sound directly^34^, possibly contributing to audiovisual multisensory integration. The source of increased auditory response strength after dark rearing is still unknown. One hypothesis is that exuberant axons from the auditory cortex that are laid out at birth are not pruned away under the influence of feedforward visual input and remain in place into adulthood^48^. This is supported by anatomical tracing studies in animals^50,51^ and human fMRI studies^52^. However, there is also anatomical evidence that auditory subcortical input might drive the visual cortex after lack of early visual experience^31,51,53^. Future experiments using functional imaging targeted to the dorsal stream neurons in combination with auditory cortical or thalamic silencing might provide a clear answer.

On the neural population level we find a clear shrinkage of the orientation manifold, accompanied by a reduced orientation decoding accuracy in specifically A/RL/AL after dark rearing. This difference seems to disappear after prolonged exposure to light in adulthood. Early visual experience is thus required to develop a neural manifold in the dorsal stream-like regions A/RL/AL that allows accurate decoding of oriented moving bars. Exposure to light in adulthood seemingly recovers this decoding impairment, which could suggest that the adult brain remains plastic. Although lower number of animals compared to the first imaging session could bias these results.

In summary, a lack of early visual experience increases auditory at the cost of visual processing in the mouse visual cortex, particularly in the dorsal stream. This could enhance dorsal stream-specific auditory processing but hamper restoration of visual tuning in adulthood.

## MATERIALS AND METHODS

### Animals

All experiments were approved by the institutional animal care and use committee of the Royal Netherlands Academy of Arts and Sciences under Central Committee Animal experiments (CCD) license AVD80100202215934, Academy of Arts and Sciences. We used X male and X female mice in the dark reared group and X male and X female mice in the normally reared group. All mice were F1 offspring from female CBA/JRj (Janvier labs, https://janvier-labs.com/) crossed to male Vipr2-Cre^-/-^ (Jackson Laboratories, https://www.jaxmice.jax.org/, strain 031332)^54^. Mice were group housed before surgery. After surgery, they were either individually or group housed with *ad libitum* access to food and water under a 12-hour reversed day/night cycle or constant darkness. All mice had access to a running wheel in their home cage. Extra care was taken to ensure dark reared animals remained in absolute darkness up until the first recording. Daily health checks and habituation to the recording setup were done under infrared light (850 nm) using custom infrared binoculars. During surgery the eyes were covered using custom light-protective caps. All experiments were performed in the dark phase of normally reared animals.

### Viral injection and cranial window surgery

Mice were anesthetized with isoflurane (5% induction, 1.5 - 2% maintenance in oxygen). Body temperature was maintained at a stable 37 °C with a euthermic pad, and the eyes were covered with light-protective caps filled with Ophtosan eye ointment to prevent them from drying out. Metacam (5 mg/kg *s.c.*), Temgesic (0.1 mg/kg *s.c.*), Xylocaine gel (on the periosteum) and Rimadyl (0.06 mg/mL, post-operative day 1 to 3 in drinking water) were provided for analgesia and Dexamethasone (4-8 mg/kg *s.c.*) for anti-inflammation and to reduce brain swelling. First, the skin and periosteum were removed above the skull. Then, eight holes were drilled in the skull spread across the visual cortex using a dental drill. A mixture (1:1) of AAV1-hDlx-dlox-mCyRFP1(rev)-dlox (0.67 x 10^12^ titer, VVF Zurich, v313-1) and AAV9-CaMKIIα-jGCaMP8m (0.67 x 10^12^ titer, VVF Zurich, v630-9)^55^ was injected at 400 and 200 µm below the cortical surface across 8 injections of 9.2 nL at 46 nL/s. A craniotomy was drilled above the visual cortex and a cranial window (stacked 4, 4 and 5 mm diameter coverslips) was carefully lowered in the craniotomy. The window and a custom metal head ring were attached to the skull using dental cement. Mice were allowed to recover from surgery for a minimum of one week, after which they were habituated to head fixation in the recording setup until they comfortably started running.

### Wide-field calcium imaging

Population receptive field mapping and field sign analysis using the wide-field macroscope was done as described previously^39^. In short, the complete 4 mm diameter window was imaged using a wide-field fluorescence macroscope (Axio Zoom.V16 Zeiss/Caenotec-Prof. Ralf Schnabel). Images were captured by a high-speed sCMOS camera (pro.edge 5.5) at 20 Hz, 1600 x 1600 pixels with 50 ms exposure time and recorded with Encephalos software (Caenotec-Prof. Ralf Schnabel) after which they were downsampled to 800 x 800 pixels for analysis.

### Two-photon calcium imaging

Two-photon calcium imaging was performed on a 2P microscope (Neurolabware) equipped with a Ti-sapphire laser (Mai-Tai ‘Deepsee’, Spectraphysics) running at 920 nm, using a 16x, 0.8 NA water immersion objective (Nikon) at 1.0x to 1.6x digital zoom. Imaging was controlled by Scanbox software (Neurolabware) running on Matlab at a frame rate of approximately 15.5 Hz. Mice were head fixed and free to run on a wheel. Running speed was recorded using a rotary encoder running on an Arduino and processed in Matlab.

### Visual stimuli

#### Wide-field calcium imaging

Mice were positioned centrally, 14 cm away from a 122 x 68 cm LCD screen (iiyama LE5564S-B1, 1920 x 1280 pixels at 60 Hz refresh rate) covering a 144° x 86° field-of-view. Stimuli were created and presented using COGENT (developed by John Romaya at the LON at the Wellcome Department of Imaging Neuroscience) together with Matlab (Mathworks). Stimuli consisted of eccentricity-corrected angled bars at 0°, 45°, 90° or 135° with a width of 10°, filled with a checkerboard pattern on a grey background (20 cd/m2). A total of 58 stimuli were pseudo-randomly presented 10 times for 0.5 s with an inter-trial-interval consisting of a grey screen (20cd/m^2^) for 3.6 s. Images were smoothed using a Gaussian filter with two pixels standard deviation and stored per trial. Population receptive field mapping and field sign analysis was done as described previously^37,39^.

#### Two-photon calcium imaging

Mice were positioned centrally, 15 cm away from a 24 inch gamma-corrected HD LED monitor (1920 x 1080 pixels at 60 Hz refresh rate) covering a 120° x 90° field-of-view. Stimuli were created and presented using OpenGL and Psychophysics Toolbox 3 in Matlab. Visual stimuli consisted of eccentricity-corrected full-field sinusoidal drifting gratings at a direction of 0°, 45°, 90°, 135°, 180°, 225°, 270° or 315° at 8 different contrasts logarithmically spaced between 1% and 100% (“logspace(0,2,8)” in Matlab). Each combination of direction x contrast was pseudorandomly presented for 1 s (0.05 cpd at temporal frequency of 1 Hz) with an inter-trial-interval of 2 s where a mean-luminance grey screen was presented. This was repeated 15 times.

### Auditory stimuli

For 2P calcium imaging, a single ultrasound speaker (Batsound, L400 Ultrasound speaker) was positioned at the level of the mouse at 20 cm distance. To measure frequency response areas, pure-tone stimuli were created and presented using Psychophysics Toolbox 3 and Matlab. Pure-tone stimuli were presented in open field at 12 frequencies from 1.5 kHz to 48 kHz in half-octave steps and at intensities of 50, 60, 70 and 80 dB sound pressure level (dB SPL) at 192 kHz with 24 bits using the Terratec Aurean Xfire 8.0 HD external soundcard. Peak intensity levels of background noise were around 40 dB SPL. Pure-tone stimuli were presented in pseudorandom order for 0.5 s (including a 5 ms cosine-squared rise/decay time) with a random inter-trial interval of 2 to 3 s. This was repeated 15 times. Sound intensities were calibrated using a condenser ultrasound microphone (Avisoft-Bioacoustics CM16/CMPA) placed at the position of the pinnae, in combination with the UltraSoundGate 116Hb recording interface and Avisoft-RECORDER USGH software.

### Video tracking and whisker energy analysis

Facial movement during 2P calcium imaging was recorded using an infrared camera (FLIR, Flea3 USB3 combined with a Tamron 12VM412ASIR 1/2” 4-12mm F/1.2 Infrared Manual C-Mount Lens) triggered by 2P image acquisition at around 15.5 Hz. Video files were further processed using Facemap^56^ and Matlab. Motion energy was extracted from a manually drawn region of interest around the whisker pad.

Z-scored whisker energy was calculated using the 2 to 0 seconds before tone onset per trial. The peak of the Z-scored whisker energy was extracted between 0 and 0.5 s and averaged across trials for analysis in Fig. S1. Trials were removed from pure-tone analysis if the mean Z-scored whisker energy in either 0.5 s after tone onset or offset was above 2.

### Two-photon calcium imaging analysis

Raw images were pre-processed using the SpecSeg toolbox^57^ to select regions of interest (ROIs) that correspond to the somata of neurons and extract the mean fluorescence signal. In brief, first, rigid motion correction was performed using NoRMCorre^58^, then ROIs were selected based on cross-spectral power across pixels. ROIs were manually refined, and mean signal was extracted per ROI. Then, neuropil was corrected by subtracting 70% of the average pixel value in a doughnut around the ROI. ΔF/F values were calculated by subtracting a moving baseline and then dividing by a linear fit of that moving baseline (10^th^ percentile over 500 frames).

To increase the temporal resolution of the ΔF/F trace we used a spike deconvolution algorithm (MLspike, autocalibration of parameters per neuron)^43^. Additionally, run speed was regressed out of the estimated spike times using a linear model (‘fitlm’ in Matlab). The residual estimation of the firing rate (FR) was used to calculate the ΔFR by subtracting the mean activity from 0.5 to 1 s before tone onset.

After removal of whisker movement trials, responses to pure-tone stimuli were calculated by taking the trial-averaged ΔFR 0 to 0.5 s after tone onset and offset or 0 to 0.5 s after drifting grating onset. A minimum of 8 trials per tone was required after trial removal; otherwise, the tone was excluded from further analysis. For calculation of response type and response amplitude the period (after tone onset or offset for pure tones or just after grating onset) with the largest significant response was taken from the stimuli with the strongest significant response (best frequency for pure-tone stimuli or preferred stimulus for drifting gratings). A neuron was deemed significantly responding if the ΔFR response from at least one stimulus type exceeded a trial-shuffled threshold. The threshold was set as the 0.1st percentile of a distribution obtained from 1,000 shuffles of ΔFR response amplitudes, where the deconvolved spike times were circularly shifted by a random time every shuffle.

### Population geometry analysis

Population analyses used deconvolved firing rates from 1000 randomly redrawn subsets of N = 10 neurons per experimental session and cortical area, reusing the same set of subsets across analyses. This subset size was chosen to ensure population activity for all cortical areas in each rearing condition and at each timepoint was measured in at least 2-3 recording sessions. For most analyses, firing rates were averaged over 0-0.5 s after stimulus onset.

For cosine similarity (Fig. 3B), trials from the two highest contrasts were pooled to obtain eight direction-specific population response vectors. These vectors were centered by subtracting their mean across directions, and cosine similarities were averaged across direction pairs separated by 90° or 180°.

Population spread was quantified as the squared radius,

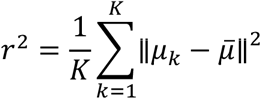

where *μ_k_* denotes a stimulus-specific mean response vector and *μ^-^* the mean of these vectors across the *K* stimuli. Squared radius was calculated in the full N=10 neuron space, using eight direction vectors for Fig. 3D and four orientation vectors, pooling opposite directions, for Fig. 3F. In Fig. 3F, radius was calculated at each imaging frame for the lowest two and highest two contrasts separately. Radius-over-time curves were averaged across neuron subsets and sessions within each mouse before taking their maximum over 0–0.5 s, which is the statistic reported in F.

### Linear discriminant analysis

Linear discriminant analysis (LDA) was used to decode four grating orientations or eight directions from the same 0–0.5 s population responses. The pooled within-class covariance was regularized using Ledoit–Wolf shrinkage estimated exclusively from training trials. Performance was evaluated using five-fold cross-validation, stratified by direction and contrast. For within-condition decoding (Fig. 3A), training and test trials were drawn from a pooled subset of trials corresponding to the two highest contrasts. For cross-condition orientation decoding, the eight contrast levels were grouped into four adjacent pairs. Classifiers were trained on four folds from a higher-contrast bin and tested on the held-out fold from the immediately lower bin, yielding transitions 4→3, 3→2 and 2→1. Accuracies were averaged across folds and neuron subsets, then across sessions within mice and equally across mice. Chance performance was 25% for orientation and 12.5% for direction.

### Dimensionality reduction

Principal component analysis (PCA) was used to visualize population geometry in Fig. 3C. Direction-specific response vectors were calculated from the two highest contrasts for D and separately from the lower four and upper four contrasts for H. Within each of the five folds, training and held-out configurations were centered across directions. Training configurations were aligned to an area-specific reference using orthogonal Procrustes transformations, which preserve distances without rescaling, and the fitted transformations were applied to held-out configurations. For H, both contrast pools were aligned jointly. Held-out coordinates were averaged across folds and neuron subsets, sessions within mice, and mice. The resulting group configurations were registered across areas, and a shared three-component PCA was fitted separately for D and H across all included areas and conditions. In H, opposite-direction coordinates were averaged after projection to display four orientation points. PCA was used to provide a descriptive qualitative visualization; quantitative geometry was measured in the full N=10 neural activity space before alignment, averaging and projection.

### Statistical analysis

Statistical details are written in the figure legends. For two-way ANOVAs and repeated measures ANOVAs the data were assumed to be normally distributed. Repeated measures two-way ANOVAs were Geisser-Greenhouse corrected. Data are plotted as mean ± SEM, except for LDA data which is plotted as mean ± SD. ANOVAs and Fisher’s exact tests were performed in Graphpad Prism 10 and Mann-Whitney U tests and permutation tests in Matlab (R2023b).

For population-level analyses, data are presented as mouse means with 95% bootstrap confidence intervals. Effects of rearing, cortical area, timepoint and the panel-specific factor were tested using rank-based nonparametric longitudinal ANOVA-type tests (nparLD package), accounting for repeated measurements within mice. NR–DR comparisons used two-sided Wilcoxon rank-sum (Mann–Whitney U) tests. Multiple comparisons were Holm-corrected within each panel, as specified in the figure legends.

## ACKNOWLEDGEMENTS

We wish to thank the Levelt lab for helpful discussion, the members of the mechatronics department for technical support and the animal caretakers of the Netherlands Institute for Neuroscience for their help. We thank Vasileios Patsourakos, Maaike van der Aa and Ayana Kemble for help with habituation of the mice and Chris van der Togt, Matthew W. Self, Pieter W. Roelfsema and Gerard Borst for technical support, data analysis and data management support. This project received funding from the European Union’s Horizon 2020 Research and Innovation Program under grant agreement nos. 785907 (HBP SGA2, CL & PR) and 945539 (HBP SGA3, CL & PR) and the Dutch Research Council (FlagEra SoundSight, 680-91-320, CL). The authors declare that they have no competing interests.

## SUPPLEMENTARY FIGURES

**Supplementary figure 1.**
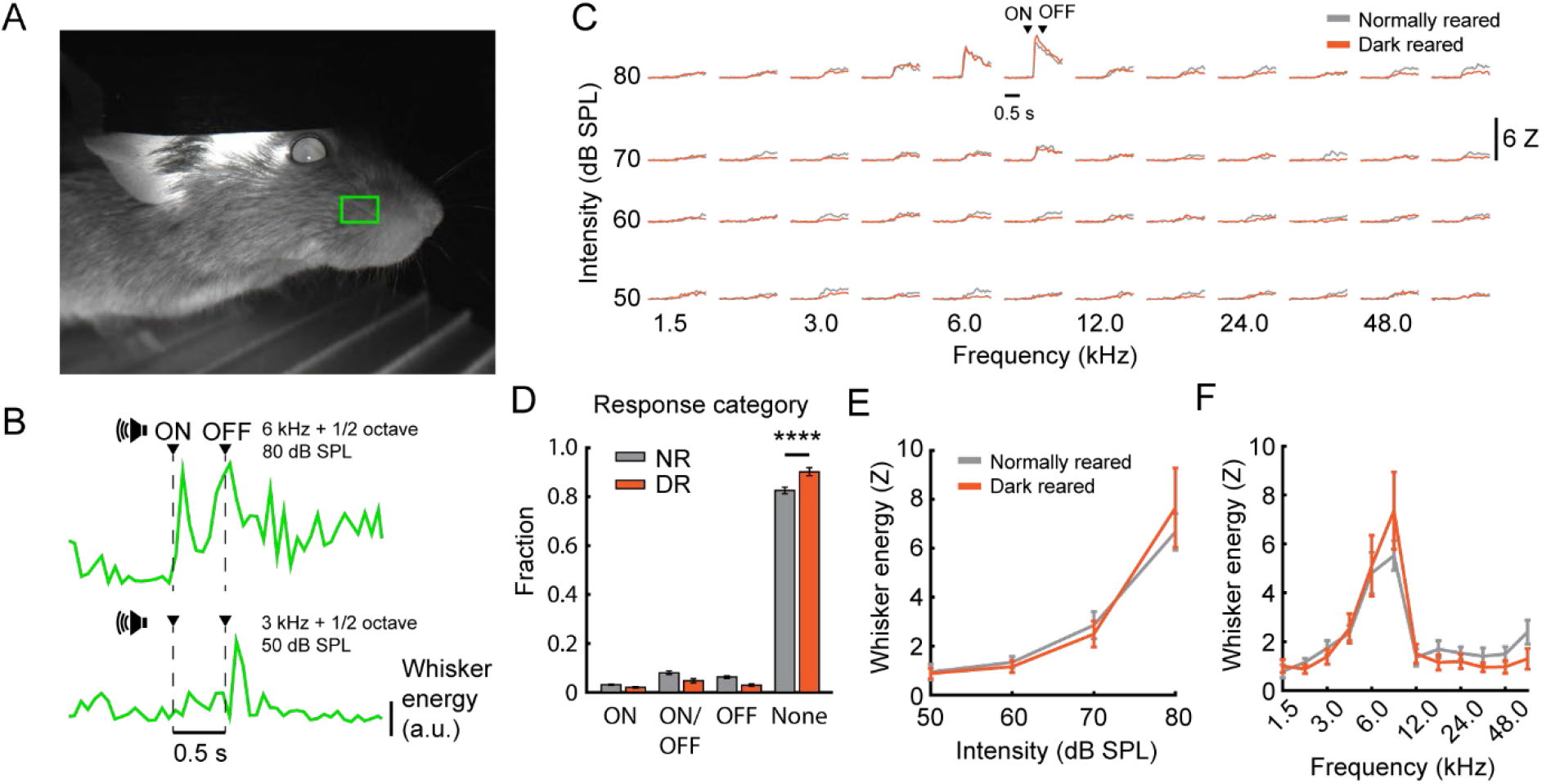
Tone intensity-dependent and frequency-dependent increase in whisker movement. (A) Example movie image from the face of a mouse in darkness with only infrared light. Green square, whisker region. (B) Example trials with whisker energy (from the green square in (A)) in response to a tone with an ON/OFF response (top) and OFF response (bottom). (C) Session-averaged frequency response area of the whisker energy. Whisker energy is z-scored to 1-0 s before tone onset. (D) Fraction of response categories. ON, average whisker energy (z-scored) between 0 to 0.5 s above 2 during this period only. ON/OFF, average whisker energy (z-scored) between 0 s to 0.5 s and 0.5 s to 1.0 s above 2. OFF, average whisker energy (Z-scored) between 0.5 s to 1.0 s above 2 during this period only, None, average whisker energy (z-scored) between 0 s to 0.5 s or 0.5 s to 1.0 s below 2. Two-way ANOVA with Dunn-Sidak multiple comparison correction, Category x rearing, F(3,168) = 15.42, p < 0.0001, NR None vs DR None, p < 0.0001. Data is session averaged over trials. Normally reared, 19 sessions, Dark reared, 25 sessions. (E) Whisker energy (z-scored) per tone intensity. Tone intensities were used across the strongest frequency response. Two-way repeated measures ANOVA. Intensity, F(2.249,88.33) = 23.49, p < 0.0001, rearing, F(1,40) = 3.273e-005, p = 0.9955, intensity x rearing, F(11,432) = 1.169, p = 0.3063. (F) Whisker energy (z-scored) per frequency. Tone frequencies were used across the strongest intensity response. Frequency, F(1.419,50.14) = 33.50, p < 0.0001, rearing, F(1,36) = 0.2513, p = 0.6192, frequency x rearing, F(3,106) = 0.7808, p = 0.5072.

## Notes

### Competing Interest Statement

The authors have declared no competing interest.

### Summary of Updates

Figure 3 updated on effect of dark rearing on population coding of visual stimuli.

